# Metastability indexes global network effects post brain stimulation

**DOI:** 10.1101/2023.11.23.568409

**Authors:** Rishabh Bapat, Anagh Pathak, Arpan Banerjee

## Abstract

Several studies have shown that coordination among neural ensembles is a key to understand human cognition. A well charted path is to identify coordination states associated with cognitive functions from spectral changes in the oscillations of EEG or MEG. A growing number of studies suggest that the tendency to switch between coordination states, sculpts the dynamic repertoire of the brain and can be indexed by a measure known as metastability. In this article, we characterize perturbations in the metastability of global brain network dynamics following Transcranial Magnetic Stimulation that could quantify the duration for which information processing is altered, thus, allowing researchers to understand the network effects of brain stimulation, standardise stimulation protocols and design experimental tasks. We demonstrate the effect empirically using publicly available datasets and use a digital twin (a whole brain connectome model) to understand the dynamic principles that generate such observations. We observed a significant reduction in metastability, concurrent with an increase in coherence following single-pulse TMS reflecting the existence of a window where neural coordination is altered. The reduction in complexity was validated by an additional measure based on the Lempel-Ziv complexity of microstate labelled EEG data. Interestingly, higher frequencies in the EEG signal showed faster recovery in metastability than lower frequencies. The digital twin shed light on how the phase resetting introduced by the single-pulse TMS in local cortical networks can propagate globally across the whole brain and give rise to changes in metastability and coherence.

## Introduction

Processing the complex dynamic environment around us requires flexible exploration of neural coordination states that helps in brain function (Deco and Kringelbach, 2016). The ability to switch between coordination states is driven by the tendency of the dynamical system to traverse through multiple attractors (Haken, 1983; Bressler, 2002; Kelso and Zanone, 2002; Tognoli and Kelso, 2014), quantitatively captured by a mathematical measure, metastability (Deco and Kringelbach, 2016; Tognoli and Kelso, 2014). Reflecting the fundamental role of metastability, modelling studies have found it to be a signature of brain’s dynamic core (Deco et al., 2017) and maximized in the resting state (Hellyer et al., 2014; Deco et al., 2017; Naskar et al., 2021; Saha et al., 2023). A plethora of studies further validate this claim by showing changes in metastability to accompany altered or disordered states of consciousness. Metastability is shown to be reduced during loss of consciousness (Cavanna et al., 2018; Jobst et al., 2017), following traumatic brain injury (Hellyer et al., 2015) and in Alzheimer’s disease (Córdova-Palomera et al., 2017). Interestingly, metastability is found to be higher among schizophrenics (Lee et al., 2018) and following the use of psychedelic drugs (Carhart-Harris et al., 2014; Lord et al., 2019). Some of these studies also indicate changes in cognitive flexibility caused by reduction in metastability (Córdova-Palomera et al., 2017; Hellyer et al., 2015). Are these changes in metastability idiosyncratic or do they arise from a principled organization of brain network dynamics? Answering this question requires hypothesis driven empirical observation followed up with theoretical understanding of whole-brain network dynamics.

Transcranial Magnetic Stimulation (TMS) is known to cause a phase-reset within the stimulated region (Kawasaki et al., 2014; Pellicciari et al., 2017). This can effectively force the underlying neural to get into a coordinated state at least transiently. From a dynamical systems standpoint, getting into an attractor state will make a high dimensional system low-dimensional and thus lead to a reduction in metastability (Pillai and Jirsa, 2017). Thus, even a single-pulse TMS can lower the metastability of brain dynamics. Hence, metastability might also be used as an index of neural dynamics to describe the effects of a given TMS protocol. Using metastability to contextualise the effects of such stimulation could help explain the variability pervasive in TMS research and help optimise therapeutic TMS protocols for disorders known to have altered metastability. Once, perfected with TMS, similar effects can be studies for more emerging methods of non-invasive brain stimulation such as transcranial Direct Current Stimulation (tDCS) and transcranial Alternating Current Stimulation (tACS).

In the present article we test the hypothesis that metastability which is typically associated with the resting state is reduced in a time-window that is time-locked to the onset of TMS pulse. The recovery to pre-stimulus levels of metastability will index the temporal window over which the network is perturbed by the stimulation and may vary with the oscillation frequency. Secondly, we illustrate that using a digital twin - a whole-brain network of phase-coupled Kuramoto oscillators connected with a bio-physically realistic connection topology (Cabral et al., 2011; Jirsa et al., 2017) - can shed light on the dynamic principles that are key to such empirical findings. Taken together, the empirical findings in tandem with the theoretical approach enhance our understanding of systems level neural mechanisms that unfold following non-invasive brain stimulation.

## 2 Methods

### 2.1 Data Collection

The data used in this study was obtained from OpenNeuro https://openneuro.org/. One dataset had 20 participants (Hussain, 2019) (Dataset 1) while the other had 13 (Pavon et al., 2022)(Dataset 2). 300 seconds of resting state, eyes open EEG were recorded for estimation of Resting Motor Threshold (RMT) using either a 30 channel electrode array and a 5kHz sampling rate (Dataset 1) or a 63 channel electrode array with a 20kHz sampling rate (Dataset 2). Single pulse, monophasic TMS was delivered to the right Primary Motor Cortex (rPMC) at 100%, 110% and 120% of RMT every 5 seconds. 600 trials at 120% of the RMT were conducted in dataset 1 with 75 trials at 110% and 100% of the RMT in the dataset 2. EEG recordings were taken simultaneously. The coil noise was not masked, but participants were provided with earplugs to reduce the disturbance.

## 2.2 Preprocessing

The resting state EEG and TMS-EEG data were pre-processed prior to analysis. The resting state EEG data were processed using custom MATLAB code utilizing a combination of Artifact Subspace Reconstruction (Plechawska-Wojcik et al., 2019) and Multiple Artifact Rejection Algorithm (MARA) (Winkler et al., 2011) based independent component rejection. The TMS-EEG data were preprocessed using the Automated Artifact Reject for Single Pulse TMS Data (ARTIST) pipeline (Wu et al., 2018). All data was downsampled to 1kHz and bandpass filtered between 1 and 100 Hz prior to analysis.

### 2.3 Measures of Metastability

#### 2.3.1 The Standard Deviation of the Kuramoto Order Parameter

Two measures of metastability were applied to the preprocessed EEG time series (Figure 1). The first measure was based on the standard deviation of the Kuramoto Order Parameter (KOP) based on the work of Yoshiki Kuramoto Kuramoto (1984). The KOP is a measure of synchrony in a network of oscillators and is calculated as the real part of the normalized vector sum of individual phases from each nodes,

**Figure 1.**
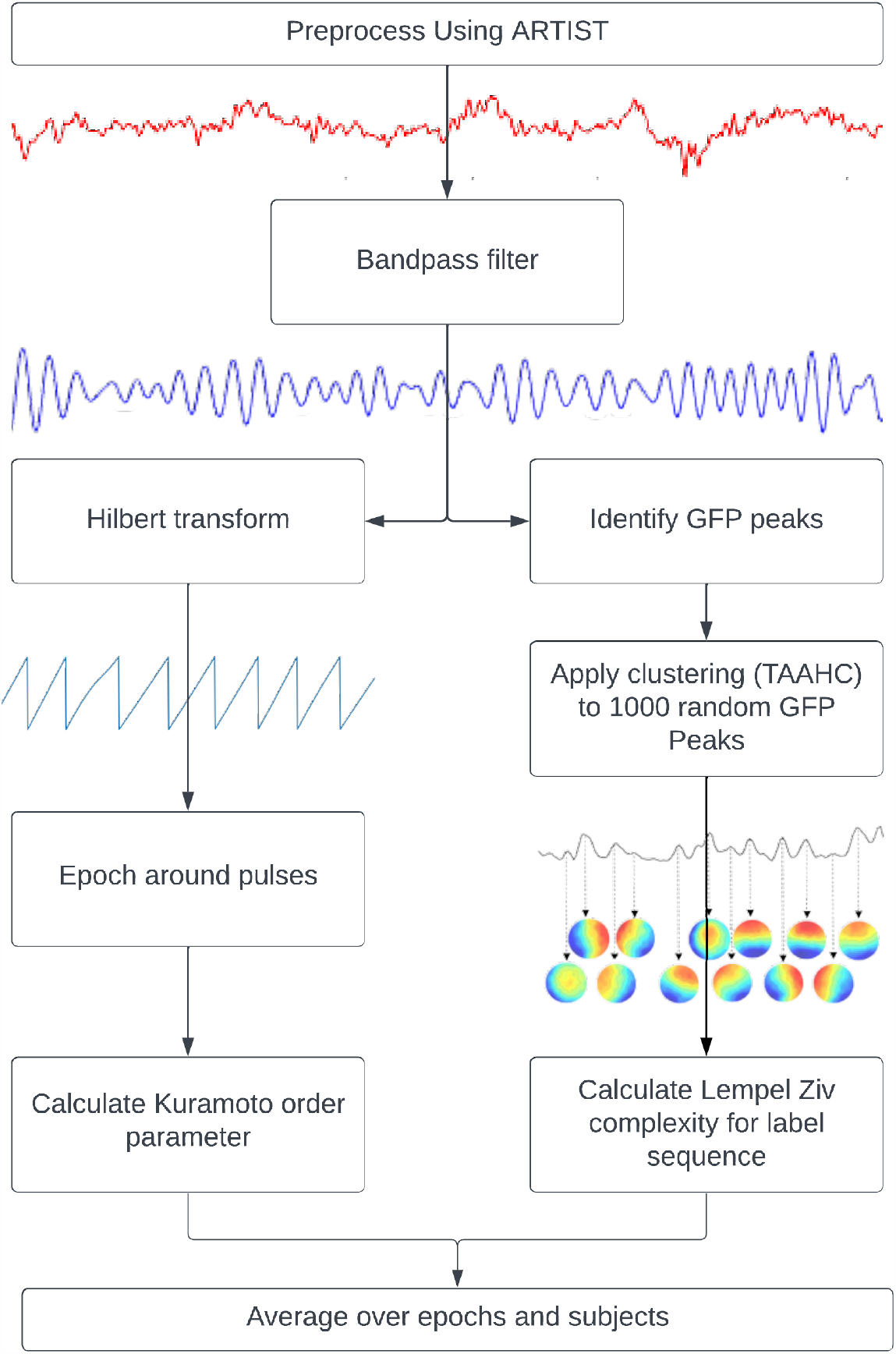
Pipelines used to calculate metastability. Kuramoto Order Parameter (KOP) on EEG data (left) and the Lempel-Ziv Complexity (LZC) on microstates (right).

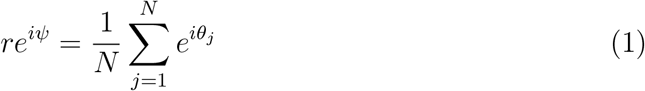

where ‘*N* ‘is the number of oscillators, ‘*θ*_*j*_’ is the phase of the *j*th oscillator, ‘*r*’ is the Kuramoto order parameter and *ψ* is the phase angle following the vector sum of individualized phases. The KOP (equation 1) can be thought of as plotting the phase of each oscillator as a point on a unit circle, and then taking the magnitude of their resultant vector. Its value is 1 for a completely synchronised system and 0 for a desynchronised system (with infinite nodes). Its standard deviation indexes the ability of the system to deviate from stable states and thus can be used as a proxy measure of metastability. For this measure, the TMS EEG data were bandpass filtered within narrow frequency bands (for example, 8 to 12 Hz), after which the instantaneous phase was extracted via Hilbert transformation. The KOP was then calculated for each time point. The standard deviation of the KOP was calculated in a sliding window of 50 ms and was used as a dynamic measure of metastability. The metastability time series was then averaged across epochs and participants to yield the final results.

The choice of which group of channels to include for computation of metastability is an important one. Since changes to global coherence across all channels are relatively less intense, and stimulation with TMS has prominent local effects, channel groupings derived from an algorithm were analyzed in addition to global metastability. This algorithm aimed to capture local effects on synchrony and grouped channels together based on the similarity of their TMS evoked potential. First the evoked response to TMS was calculated in each stimulation intensity by averaging across epochs and subjects. Then the time point with the highest Global Field Power (GFP), meaning the highest spatial standard deviation, was selected. At this time point, the activity of each channel was subtracted from the mean of all channels. The channels were then sorted in ascending order of their mean differences, and the top and bottom tertiles were assigned to separate groups. This algorithm is a quantitatively rigorous way to select channels as opposed to visually inspecting the evoked response, finding the point at which the channels activity is the most varied, and sorting the most extreme channels into their own groups.

#### 2.3.2 Microstate Sequence Complexity

An additional measure of metastability used in this study is based on microstate sequence complexity. Microstates are quasi-stable spatial activity patterns that are derived from EEG data using clustering algorithms. The original recording can then be backfitted to the most similar microstate at each point in time and be analysed as a sequence of microstates (Lehmann, 1971). The quasi-stable nature of these states is reminiscent of the dwell and escape tendencies seen in metastable dynamics while the microstates themselves are related to the coordination states. The central idea behind this measure is that metastable coordination dynamics will produce a non-repetitive microstate sequence. This can be quantified for the microstate sequence using measures of ‘complexity’.

‘Complexity’ refers to the unpredictability of a signal and is quantified here by Lempel-Ziv Complexity (LZC). The LZC of a string is the minimum number of unique sub-strings that can be repeated and combined to reproduce the original. It increases with the unpredictability and length of a string (Lempel and Ziv, 1986). In order to compute this measure, microstates were derived from GFP peaks in resting state data that was bandpassed to alpha (8-12 Hz). The clustering was done using Topographical Atomise and Agglomerate Hierarchical Clustering. In this method initially, each time point is its own microstate. Through iteratively eliminating (atomising) and redistributing (agglomerating) the worst microstate, based on the sum of correlations between the microstate and its members (Correlation Sum), the number of microstates is reduced to two (a preset minimum) thereafter, an optimal number of microstates can be selected (Khanna et al., 2014; Poulsen et al., 2018). The number of microstates for each participant were selected using the Krzanowski-Lai criterion (Krzanowski and Lai, 1988) applied to the Correlation Sum. This method involves identifying the point past which the Correlation Sum plateaus with respect to the number of microstates. The microstate decomposition was performed for each participant individually. The microstates for each participant were then back fitted to their resting state data, and fitted to their TMS EEG data. After this, the LZC was calculated in a 100 ms sliding window and averaged across epochs and subjects before being compared between resting state and TMS EEG recordings. This short window was considered suitable since good temporal resolution was required for this analysis and previous literature has shown that microstate sequences show scale free dynamics (Van De Ville et al., 2010).

To test the changes in metastability for significance, two 250 ms windows of time were defined, before and after the pulse. Then, the epoch-averaged measures were averaged in the windows of time, yielding a pre and post pulse metastability value for each participant. Given that the sample size was less than 30 and the results of the Shapiro-Wilk test of normality were inconsistent, normality could not be safely assumed. Thus the difference between the two lists of metastability measures was tested for significance using Wilcoxon’s Signed Rank test. This is a non-parametric test suitable for testing dependent samples. The significance threshold was set to 0.05, and p values below this were used to indicate a significant difference. The test was carried out in a one-tailed manner, with the direction being dependent on the effect in question.

### 2.4 Computational Modelling

In order to provide mechanistic insights into metastability modulation post-TMS, a computational model was implemented similar to that used by Pathak et al. (2022). The brain was reduced to a system of 90 coupled oscillators with activity at each oscillator being generated based on the Kuramoto model,

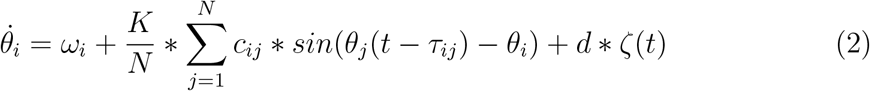

Here, the derivative of the phase of each oscillator 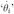 is calculated using its intrinsic frequency ‘*ω*_*i*_’, a delayed coupling term indexed by mutual coupling with oscillator of another node with a time delay *τ*_*ij*_ and coupling strength (*c*_*ij*_) and noise (*d ζ*(*t*)). Euler integration is then used to generate a phase time series for the network. The intrinsic frequencies of the network were assigned based on anatomical node strength, as described in Gollo et al. (2017). The adjacency matrix was derived from probabilistic tractography applied to diffusion MRI data and were obtained from the public repository associated with Cabral et al. (2014). The connectivity was reduced to a 90 by 90 matrix based on the Automated Anatomical Labelling parcellation scheme (Tzourio-Mazoyer et al., 2002) and averaged across subjects as in Cabral et al. (2014). The delays were obtained by scaling cortical distances by conduction velocity. Noise was sampled from a normal distribution with a mean of zero and a standard deviation of 1, it was scaled by a factor of 3. The scaling factors for coupling *c*_*ij*_ and the conduction velocity that scales *τ*_*ij*_ were chosen such that the model showed metastable dynamics, as approximated by the Kuramoto Order Parameter. The TMS pulse was modelled as a phase reset for the entire network. A subgroup of oscillators with similar intrinsic frequencies to the oscillator corresponding to the right primary motor cortex were chosen for further analysis. This subgroup is analogous to the subgroups derived from the empirical data. Metastability was then computed in a 500 millisecond sliding window based on the KOP. This analysis was repeated for frequency distributions with maximum frequencies of 5 Hz, 15 Hz, 25 Hz and 35 Hz to observe the effects of intrinsic frequency on the dynamics. The dispersion of the frequency distribution and the other parameters of the model were kept consistent. The metastability and coherence time series were produced for 20 random number generator seeds and averaged.

## 3 Results

### 3.1 Localization of TMS Induced Neural Coordination

Our first goal was to identify the subgroup of EEG sensors where the relationship between phase synchrony and metastability is most evident following the line of reasoning outlaid in section 2.3.1. We extracted the TMS pulse evoked event related potential (ERP) and undertake a channel grouping algorithm described in section 2.3.1 (Figure 2a). The channel grouping analysis extracted two distinct groups of sensors, a fronto-central and a temporo-occipital cluster. Thus, the groups appear to be clustered either around the point of stimulation (right Primary Motor Cortex) and reflected the shape of the induced electrical field.

**Figure 2.**
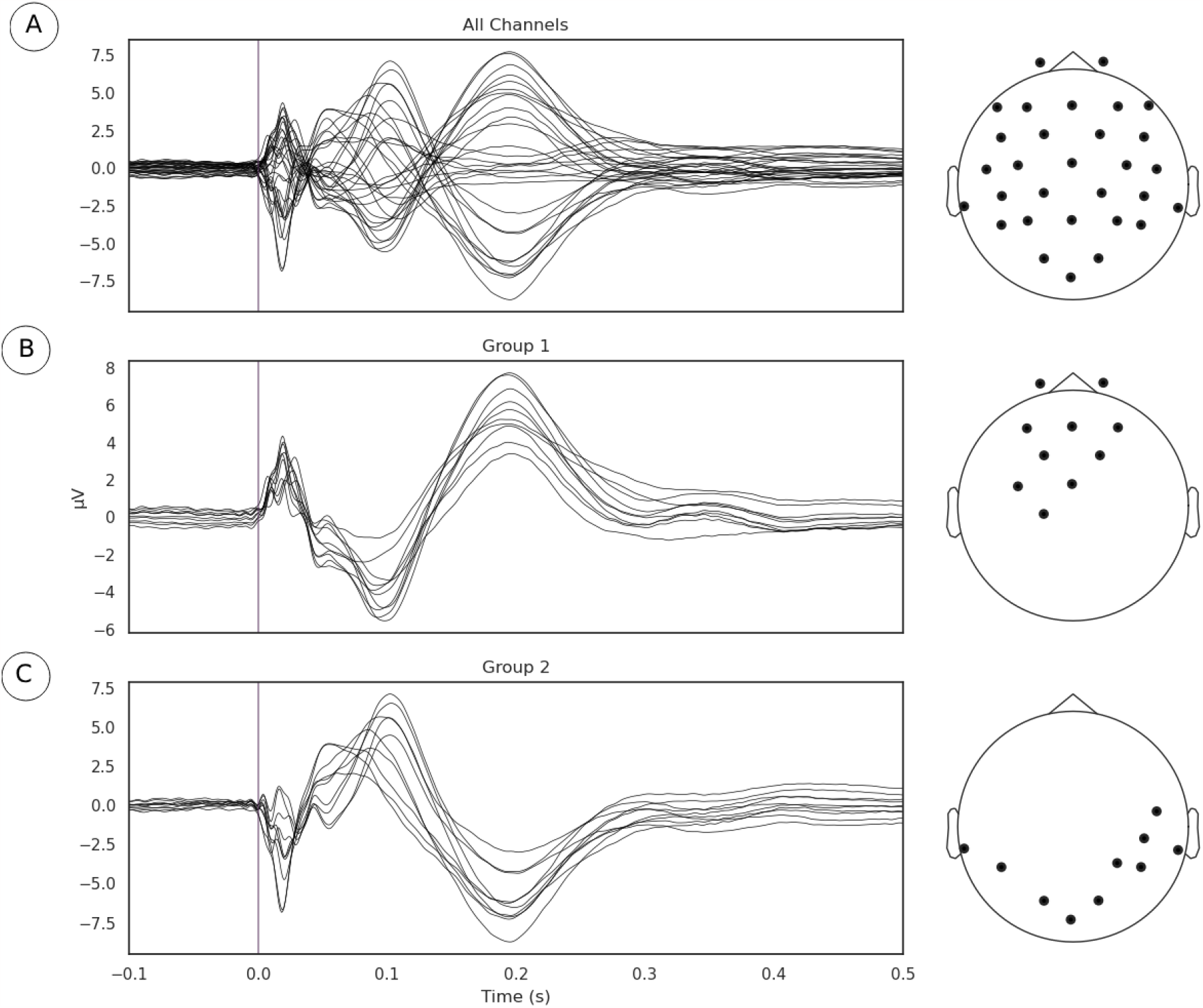
A) Evoked potentials, time locked to onset of TMS pulse. B) and C) Channel groupings or the regions of interest for examination derived from measurements of local synchrony.

### 3.2 Metastability in Pre and Post TMS Periods

Metastability was computed using two measures to demonstrate that the phenomena are robust. One measure was based on the Kuramoto order parameter (KOP), while the other used Lempel-Ziv complexity based on microstates. While the KOP measures the levels of synchrony in a select group of sensors, the standard deviation of the KOP indexes metastability (Deco et al., 2017; Pathak et al., 2022). The LZC measure is introduced in this study as an alternative measure of metastability (see details in Section 2.3.2).

The Kuramoto Order Parameter (KOP) was computed for the whole-brain scenario (all channels) and on the sub-groups of channels (fronto-central and occipito-temporal) to identify differences in the effect at the global and local level. Among all 3 groups, single pulse TMS causes a reduction of 10 to 40% in the standard deviation of the KOP calculated in different frequency bands (see Section 2.3.1). Higher frequencies (alpha, beta and gamma) recover quickly (within 200 ms) while lower frequencies (delta and theta) recover slowly (within 400 ms). Interestingly, in all frequency bands, a 10% increase prior to the pulse is seen, while in the alpha and theta bands subsequent recovery 10 to 15% past baseline levels was also observed. At a global level these changes were accompanied by a 10% decrease in KOP, however, for the subgroups, a sharp increase in KOP was observed concomitant with the same changes to metastability. These effects were replicated across all stimulation intensities, 120%, 110% and 120% of Resting Motor Threshold (RMT). The results for group 1 (fronto-central channels) and 120% RMT are plotted in Figure 3 and Table 1 details the significance of effects across different frequency bands. The same figures for the other channel groups and stimulation intensities can be found in the Supplementary Material.

**Table 1.** Test statistics and p values for one-tailed Wilcoxon’s Signed Rank tests conducted on the Kuramoto Order Parameter measure (KOP).

|  | Delta | Theta | Alpha | Beta | Gamma | All |
| --- | --- | --- | --- | --- | --- | --- |
| Decrease in KOP after pulse | 200<br>($p < 0.001$) | 210<br>($p < 0.001$) | 88<br>($p = 0.73$) | 146<br>($p = 0.06$) | 202<br>($p < 0.001$) | 205<br>($p < 0.001$) |
| Increase in KOP before pulse | 6<br>($p < 0.001$) | 2<br>($p < 0.001$) | 21<br>($p < 0.001$) | 26<br>($p < 0.001$) | 13<br>($p < 0.001$) | 149<br>($p = 0.951$) |
| Increase in KOP after pulse | 162<br>($p = 0.985$) | 55<br>($p = 0.031$) | 17<br>($p < 0.001$) | 48<br>($p = 0.016$) | 82<br>($p = 0.204$) | 197<br>($p = 0.999$) |

**Figure 3.**
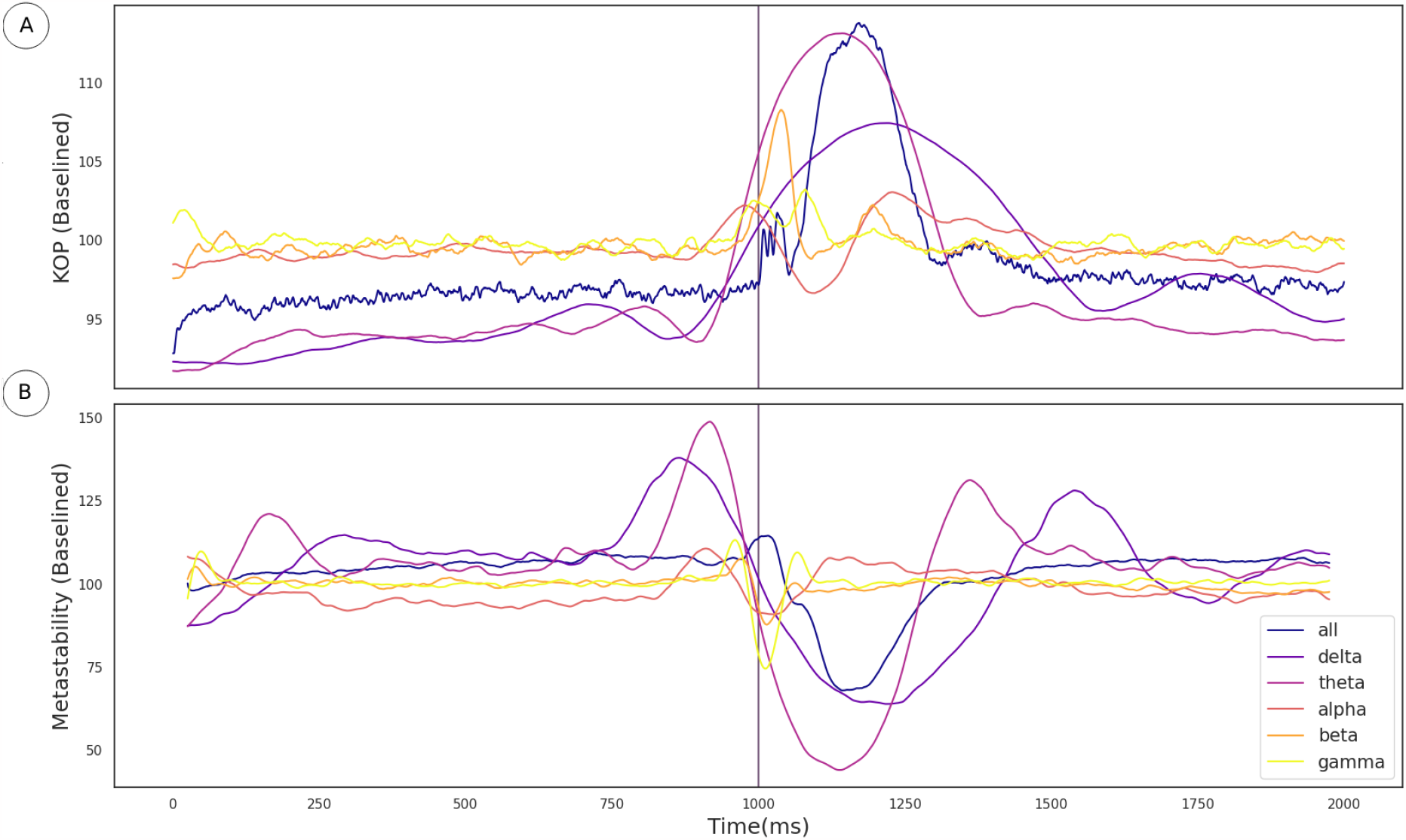
A) The effect of the TMS pulse on the Kuramoto Order Parameter. B) The effect of the TMS pulse on Metastability calculated in a sliding window. The TMS pulse is indicated by the vertical purple line. Both metastability and coherence are plotted as a percentage of a baseline value calculated as the mean between 525 and 1525 ms. Results are plotted in unique colors for each frequency band as per the legend. These results pertain to the fronto-central electrode group and the 120% RMT stimulation condition as described in Section 3.2.

To cross-validate the KOP measure of metastability, we calculated the Lempel-Ziv complexity of the sequence of microstates derived from resting state EEG data using a clustering algorithm (see Section 2.3.2). Microstate global explained variance was worse for the backfitted post-TMS EEG than for the resting state data but the performance was sufficient for continued analysis (0.65 to 0.60). Microstate analysis showed increased variation in microstate duration, and polarization of transition probabilities in the post-TMS data (Figure 4a). Note that the transition probabilities shown pertain to a single subject, since the subject specific microstate fitting made an averaging procedure unviable. However, the effect remains consistent across subjects. The LZC of the microstate sequence increases prior to the TMS pulse (test statistic = 22, p = 0.002), is reduced following TMS stimulation (test statistic = 138, p value = 0.01) and recovers to baseline levels within 200 ms (Figure 4b).

**Figure 4.**
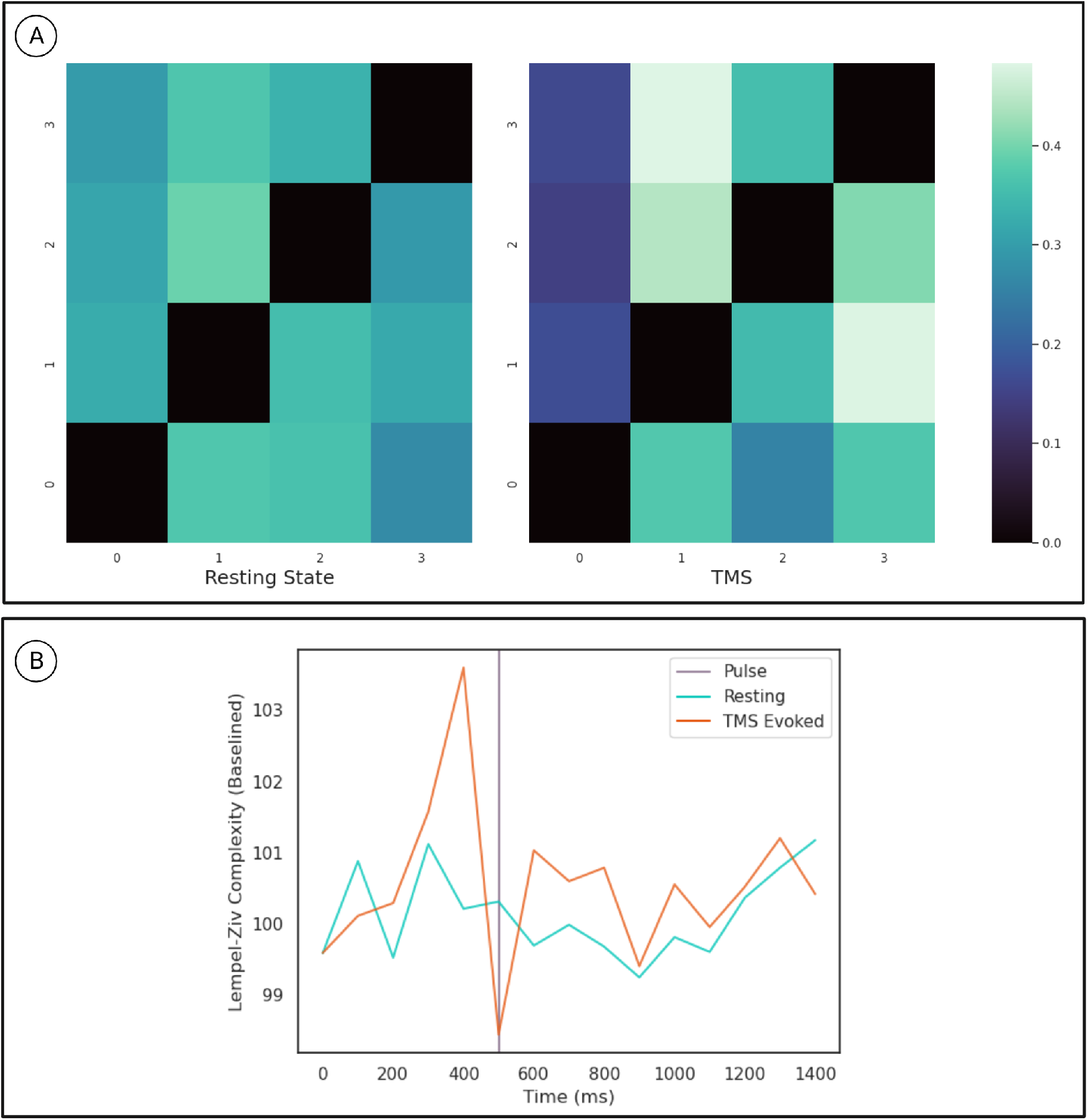
A) Microstate Transition Probabilities. The probability of transitioning from the row microstate to the column microstate is given by the color of the cell as defined by the color map on the right. The left heat-map depicts transition probabilities for 1000ms of resting state data from a given participant. While the right heat-map depicts the same for an equivalent length of time following TMS stimulation. B) Results for the Lempel-Ziv Complexity Measure. The orange line is LZC in the TMS stimulated condition while the blue line is LZC in the resting state. Complexity was calculated in 100 ms bins and averaged across subjects and epochs. The resting state data was sub-sampled and averaged in the same way. Results are plotted as a percentage of the mean LZC across the first 3 bins.

The transition probabilities for the microstates were initially relatively consistent, with each microstate having a roughly even chance of transitioning to any other microstate. In the period after the TMS pulse for which metastability is reduced, transition probabilities increase along some columns and decrease along others. Since the columns indicate the probability of another microstate transitioning into a given microstate, this change reflects the repeated consolidation of dynamics to the same pattern(s) of activity.

### 3.3 Understanding Post TMS Modulation of Metastability Using a Digital Twin

Numerical integration of the Kuramoto phase oscillators connected via bio-physically realistic coupling parameters (*c*_*ij*_, *τ*_*ij*_) (Figure 5a) was conducted using Euler integration in customized Python scripts. We identified the values of *K* and conduction velocity (that scales *τ*_*ij*_) for which the network is maximally metastable. The TMS pulse was simulated by resetting the phase of all oscillators at a given time point. The coherence and metastability was then computed from the simulated phase time series. The phase time series were simulated for multiple frequency distributions while keeping the other parameters constant (see Section 2.4).

**Figure 5.**
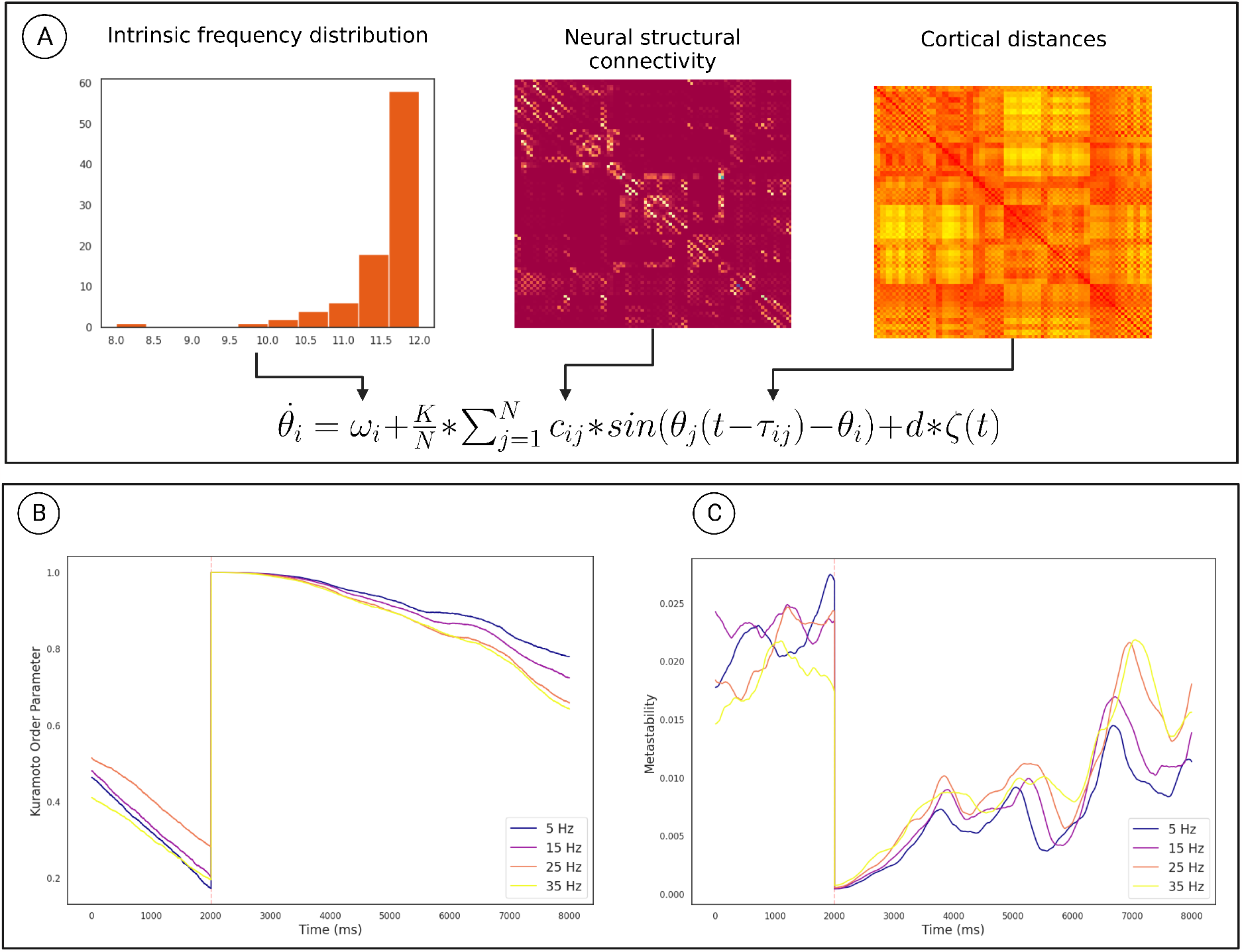
Overview of the computational model used to demonstrate the mechanism. A) How the model parameters were derived from neural connectivity. B) The effect of the phase reset on the The Kuramoto Order Parameter. c) The effect of the phase reset on the metastability calculated in a sliding window. The orange line marks the time of stimulation. The results for each frequency distribution is plotted in a unique color whose highest frequency is described by the legend. These results pertain to a subgroup of oscillators described in section 2.4.

Coherence increased following the TMS pulse followed by a reduction and recovery towards pre-TMS condition (Figure 5b). On the other hand, metastability decreases following TMS followed by a recovery towards baseline. Both of these findings were also observed in the empirical data. Furthermore, upon running the analysis for different maximum frequencies we observed that a quicker recovery of metastability occurs for higher frequency and slower recovery to baseline for lower frequency (Figure 5c).

## 4 Discussion

Metastability emerges when a delicate balance between integrative and segregative tendencies exists in a high dimensional large-scale network (Tognoli and Kelso, 2014). One can posit metastability as a winner-less competition between a set of coordination states (Deco et al., 2017). Emerging research have highlighted the usefulness of applying the measure of metastability for motor coordination (Tognoli and Kelso, 2014), lifespan ageing (Naik et al., 2017), resting state brain dynamics (Deco et al., 2017) and mental health (Deco and Kringelbach, 2016). However, the changes in brain dynamics following brain stimulation techniques such as TMS, tDCS or tACS remains to explored in detail. Since the TMS pulse may lead to a forced synchronization certain groups of oscillators, it would follow that the natural balance between them would be disrupted and the dynamics would appear less metastable. Although this effect would be highly localized and brief due to the stimulation alone, the impulse delivered to the stimulated population could propagate through the network and reset their phase dynamics as it does so. This may effectively cluster neural populations based on their collective dynamics arising out of an interaction between synaptic weights and propagation delays. The relative simplicity and stability of the resulting coordination state would manifest in reduced metastability (see Figure 6). We proved this hypothesis in the present manuscript and validated our results using two independent measures of metastability, the standard deviation of Kuramoto order parameter (KOP) and Lempel-Ziv complexity (LZC) (Figures 3 and 4). We also observed a time-scale separation in recovery of metastability from higher frequency EEG components recovering faster than lower frequencies. Digging deeper to understand the mechanistic origins of this phenomenon we employed a coupled Kuramoto oscillator network using bio-physically realistic parameters. Both findings, metastability reduction and time scale separation of the recovery trajectory were replicated by the dynamical model giving us key insights to the network interactions that unfold during TMS.

**Figure 6.**
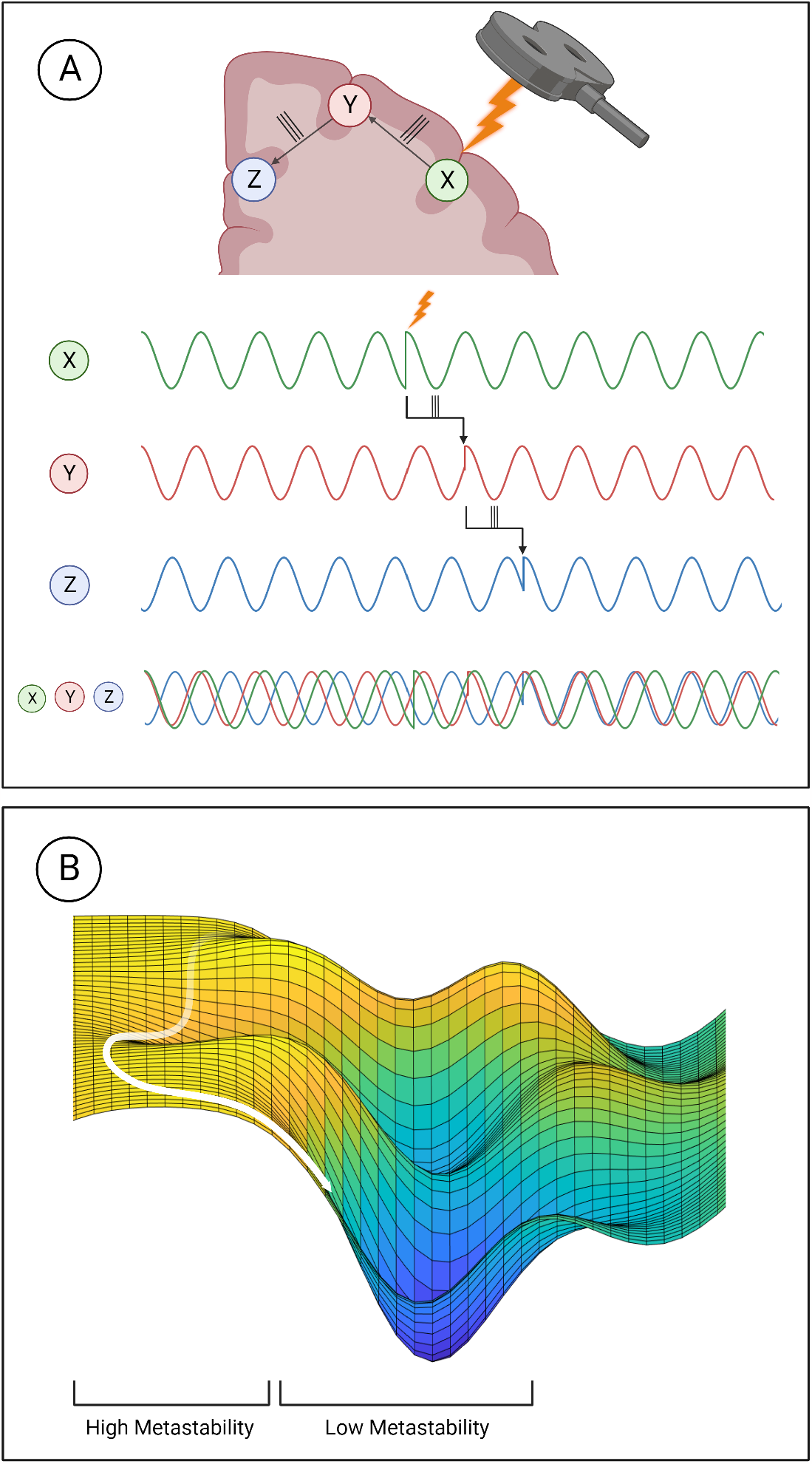
Mechanistic explanation of how TMS perturbs metastability. A) The phase locking caused by TMS can propagate through the network, increasing co-herence within clusters and decreasing the overall metastability. B) Explanation of metastability in terms of an energy landscape. The system has high metastability when it moves in and out of points of stability (wells in the energy landscape) the period of depressed metastability is analogous to a valley, where movement is more constrained.

The reduced Lempel-Ziv complexity following TMS pulse corroborates the results of the Kuramoto order parameter based analysis. Furthermore, following TMS, the microstate dynamics show increased transitions into certain microstates and reduced transitions into others. This polarisation of the microstate transition probabilities suggests that the brain repeatedly consolidates to the same coupling state under reduced metastability. This aligns with how natural dynamics would periodically start to emerge but then be interrupted as the phase reset propagates through the network with finite time delays. The results of the modelling elucidates how reduced metastability post-TMS emerge from the dynamic interactions between structural parameters such as fiber width conduction speeds. Thus our approach demonstrates how a transient phase reset can produce a measurable reduction in metastability and the dissociation of its recovery in a tonotopic organization.

The validation of metastability as a measure of coordination dynamics in short timescales is the primary contribution of this work. From a practical point of view this opens up opportunities for designing paradigms where an experimenter can choose to present task designs in a phase of reduced metastability if the effects of TMS has to be maximized. Alternatively, long term effects of TMS can be studied by presentation of stimulus at a temporal window where metastability has recovered. Furthermore, given how metastability is deranged in a wide range of neurological disorders (Cavanna et al., 2018; Hellyer et al., 2015; Córdova-Palomera et al., 2017; Lee et al., 2018), stimulation protocols aimed at treating them could show better effects if optimised based on their effect on metastability. It bears advantages over simpler measures such as coherence because of its relevance to pathology and its occurrence at multiple temporal and spatial scales. This property is exceptionally useful since it allows this measure to be applied to both EEG and MRI data, and capture changes occurring over disparate temporal and spatial scales.

An interesting and surprising result of this analysis was the increase in metastability observed prior to the pulse in the alpha and theta bands. Having thoroughly ruled out any artifactual sources of this effect, the most likely explanation was anticipation. Since hundreds of pulses were delivered during recording sessions, the distinctive ‘click’ of the TMS coil was not masked and the inter-stimulus interval was consistent, it was possible that the subjects came to anticipate the pulse and the increase in metastability reflected that mental state. To address this question, the data were examined for the presence of a Stimulus Preceding Negativity (SPN). The SPN is an ERP associated with the anticipation of an affective or physiologically arousing stimulus such as opposite sex nudes or painful electric shocks (Luck and Kappenman, 2011). In one study, an electric shock was delivered 100 - 300ms after an audio cue, and the SPN was observed in the time after the audio cue (Tanovic and Joormann, 2019). These conditions are highly reminiscent of the TMS coil click being followed by the scalp sensations of TMS. The presence of the SPN in the data (see Supplementary Information for details) and the fact that TMS can produce sensations similar to electrical stimulation, supports the idea that participants were anticipating the stimulus in the same time window as the increase in metastability. The neurophysiological mechanism for this effect and whether the increase in metastability is facilitatory or collateral to anticipation is an area of future investigation. Paired and repetitive TMS are also known to produce long term effects on coherence and excitability which could alter metastability (Bharath et al., 2023).

Another intriguing implication of this work is that metastability may change in tandem with emotional or cognitive states, as indicated by the increase during the SPN. Studying how metastability changes during cognitive tasks and in response to affective stimuli would shed light on theoretical questions of how mood can affect cognition and how dynamical properties like metastability relate to cognitive function. Finally, the global desynchronisation and local synchronisation created by TMS presents a unique lens with which to analyse it’s effects. Rather than seeing TMS as tool to excite one region, it might instead be seen as a way of linking a set of regions. Building a computational model of this phenomenon and using it to contextualise the effects of various TMS protocols is a direction of future study.

## Supporting information

Supplementary Results

## Author Contributions

RB: Design, investigation, methods, formal analysis, writing manuscript; AP: Methods, reviewing and editing manuscript; AB: Design, investigation, reviewing and editing manuscript, supervision.

## Acknowledgments

Authors acknowledge fruitful discussions with Dr Suman Saha during analysis. Authors also acknowledge the Computing Facility of NBRC for support with infrastructure.

## Data Availability Statement

The datasets analysed for this study were found on OpenNeuro https://openneuro.org/. Dataset 1 was accessed from https://openneuro.org/datasets/ds002094/versions/1.0.0, while Dataset 2 was accessed from https://openneuro.org/datasets/ds004024/versions/1.0.1. The connectivity and distance matrices were found in https://github.com/juanitacabral/NetworkModel_Toolbox.

## References

R. Bharath, Sujas Bhardwaj, R. Panda, Albert Stezin, S. Khokhar, S. Prasad, Vidhi Tyagi, N. Kamble, Keshav J. Kumar, M. Netravathi, R. Yadav, J. Annen, K. Udupa, R. Kashyap, Steven Laureys, G. Deco, and P. Pal. Single session of rtms enhances brain metastability and intrinsic ignition. 2023. doi: 10.1101/2022.08.14.503887. URL https://www.semanticscholar.org/paper/10b3f37a19c39535ba06d3d04e7c1b3760129b4b.

Steven L. Bressler. Understanding cognition through large-scale cortical networks. Current Directions in Psychological Science, 11(2):58–61, 2002. ISSN 0963-7214. doi: 10.1111/1467-8721.00168.

J. Cabral, H. Luckhoo, M. Woolrich, M. Joensson, H. Mohseni, A. Baker, M. Kringelbach, and G. Deco. Exploring mechanisms of spontaneous functional connectivity in meg: How delayed network interactions lead to structured amplitude envelopes of band-pass filtered oscillations. 2014. doi: 10.1016/j.neuroimage.2013.11.047. URL https://www.semanticscholar.org/paper/3e92ddefa41b5cb4ba3527b804d9b7acbccbed80.

Joana Cabral, Etienne Hugues, Olaf Sporns, and Gustavo Deco. Role of local network oscillations in resting-state functional connectivity. NeuroImage, 57(1): 130–139, July 2011. ISSN 1053-8119. doi: 10.1016/j.neuroimage.2011.04.010. URL https://www.sciencedirect.com/science/article/pii/S1053811911003880.

Robin Carhart-Harris, Robert Leech, Peter Hellyer, Murray Shanahan, Amanda Feilding, Enzo Tagliazucchi, Dante Chialvo, and David Nutt. The entropic brain: a theory of conscious states informed by neuroimaging research with psychedelic drugs. Frontiers in Human Neuroscience, 8, 2014. ISSN 1662-5161. doi: 10.3389/fnhum.2014.00020.

Federico Cavanna, Martina G. Vilas, Matías Palmucci, and Enzo Tagliazucchi. Dynamic functional connectivity and brain metastability during altered states of consciousness. NeuroImage, 180:383–395, October 2018. ISSN 1053-8119. doi: 10.1016/j.neuroimage.2017.09.065. URL https://www.sciencedirect.com/science/article/pii/S1053811917308133.

Aldo Córdova-Palomera, Tobias Kaufmann, Karin Persson, Dag Alnæs, Nhat Trung Doan, Torgeir Moberget, Martina Jonette Lund, Maria Lage Barca, Andreas Engvig, Anne Brækhus, Knut Engedal, Ole A. Andreassen, Geir Selbæk, and Lars T. Westlye. Disrupted global metastability and static and dynamic brain connectivity across individuals in the alzheimer’s disease continuum. Scientific Reports, 7(1):40268, January 2017. ISSN 2045-2322. doi: 10.1038/srep40268. URL https://www.nature.com/articles/srep40268. Number: 1 Publisher: Nature Publishing Group.

Gustavo Deco and Morten L. Kringelbach. Metastability and coherence: Extending the communication through coherence hypothesis using a whole-brain computational perspective. Trends in Neurosciences, 39(3):125–135, March 2016. ISSN 0166-2236. doi: 10.1016/j.tins.2016.01.001. URL https://www.sciencedirect.com/science/article/pii/S0166223616000023.

Gustavo Deco, Morten L. Kringelbach, Viktor K. Jirsa, and Petra Ritter. The dynamics of resting fluctuations in the brain: metastability and its dynamical cortical core. Scientific Reports, 7(1):3095, June 2017. ISSN 2045-2322. doi: 10.1038/s41598-017-03073-5. URL https://www.nature.com/articles/s41598-017-03073-5. Number: 1 Publisher: Nature Publishing Group.

Leonardo L. Gollo, James A. Roberts, and Luca Cocchi. Mapping how local perturbations influence systems-level brain dynamics. 160:97–112, 2017. ISSN 1053-8119. doi: 10.1016/j.neuroimage.2017.01.057.

Hermann Haken. Synergetics, 1983. ISSN 0172-7389.

Peter J. Hellyer, Murray Shanahan, Gregory Scott, Richard J. S. Wise, David J. Sharp, and Robert Leech. The control of global brain dynamics: Opposing actions of frontoparietal control and default mode networks on attention. Journal of Neuroscience, 34(2):451–461, January 2014. ISSN 0270-6474, 1529-2401. doi: 10.1523/JNEUROSCI.1853-13.2014. URL https://www.jneurosci.org/content/34/2/451. Publisher: Society for Neuroscience Section: Articles.

Peter J. Hellyer, Gregory Scott, Murray Shanahan, David J. Sharp, and Robert Leech. Cognitive flexibility through metastable neural dynamics is disrupted by damage to the structural connectome. Journal of Neuroscience, 35(24):9050–9063, June 2015. ISSN 0270-6474, 1529-2401. doi: 10.1523/JNEUROSCI.4648-14.2015. URL https://www.jneurosci.org/content/35/24/9050. Publisher: Society for Neuroscience Section: Articles.

Sara Hussain. Single-pulse open-loop tms-eeg dataset, February 2019. URL https://openneuro.org/datasets/ds002094/versions/1.0.0. Type: dataset.

Viktor Jirsa, T. Proix, Dionysios Perdikis, M. Woodman, Huifang E. Wang, J. González-Martínez, C. Bernard, C. Bénar, M. Guye, P. Chauvel, and F. Bartolomei. The virtual epileptic patient: Individualized whole-brain models of epilepsy spread. 2017. doi: 10.1016/j.neuroimage.2016.04.049. URL https://www.semanticscholar.org/paper/b2f90135f151593d4098dfa20e57fbac3e8b1e25.

Beatrice M. Jobst, Rikkert Hindriks, Helmut Laufs, Enzo Tagliazucchi, Gerald Hahn, Adrián Ponce-Alvarez, Angus B. A. Stevner, Morten L. Kringelbach, and Gustavo Deco. Increased stability and breakdown of brain effective connectivity during slow-wave sleep: Mechanistic insights from whole-brain computational modelling. Scientific Reports, 7(1):4634, July 2017. ISSN 2045-2322. doi: 10.1038/s41598-017-04522-x. URL https://www.nature.com/articles/s41598-017-04522-x. Number: 1 Publisher: Nature Publishing Group.

Masahiro Kawasaki, Yutaka Uno, Jumpei Mori, Kenji Kobata, and Keiichi Kitajo. Transcranial magnetic stimulation-induced global propagation of transient phase resetting associated with directional information flow. Frontiers in Human Neuroscience, 8, 2014. ISSN 1662-5161. doi: 10.3389/fnhum.2014.00173.

J. A. S. Kelso and P. G. Zanone. Coordination dynamics of learning and transfer across different effector systems. Journal of Experimental Psychology: Human Perception and Performance, 28(4):776–797, 2002. ISSN 1939-1277. doi: 10.1037/0096-1523.28.4.776.

Arjun Khanna, Alvaro Pascual-Leone, and Faranak Farzan. Reliability of resting-state microstate features in electroencephalography. PloS One, 9(12): e114163, 2014. ISSN 1932-6203. doi: 10.1371/journal.pone.0114163.

W. J. Krzanowski and Y. T. Lai. A criterion for determining the number of groups in a data set using sum-of-squares clustering. Biometrics, 44(1):23–34, 1988. ISSN 0006-341X. doi: 10.2307/2531893. URL https://www.jstor.org/stable/2531893.

Y. Kuramoto. Chemical oscillations, waves, and turbulence. 1984. doi: 10.1007/978-3-642-69689-3. URL https://www.semanticscholar.org/paper/f6f215bf741217b7acf66ffe301dd11b2a4f4d2b.

Won Hee Lee, Gaelle E. Doucet, Evan Leibu, and Sophia Frangou. Resting-state network connectivity and metastability predict clinical symptoms in schizophrenia. Schizophrenia Research, 201:208–216, November 2018. ISSN 0920-9964. doi: 10.1016/j.schres.2018.04.029. URL https://www.sciencedirect.com/science/article/pii/S0920996418302421.

D Lehmann. Multichannel topography of human alpha EEG fields. Electroencephalography and Clinical Neurophysiology, 31(5):439–449, November 1971. ISSN 0013-4694. doi: 10.1016/0013-4694(71)90165-9. URL https://www.sciencedirect.com/science/article/pii/0013469471901659.

A. Lempel and J. Ziv. Compression of two-dimensional data. IEEE Transactions on Information Theory, 32(1):2–8, January 1986. ISSN 1557-9654. doi: 10.1109/TIT.1986.1057132.

Louis-David Lord, Paul Expert, Selen Atasoy, Leor Roseman, Kristina Rapuano, Renaud Lambiotte, David J. Nutt, Gustavo Deco, Robin L. Carhart-Harris, Morten L. Kringelbach, and Joana Cabral. Dynamical exploration of the repertoire of brain networks at rest is modulated by psilocybin. NeuroImage, 199:127–142, October 2019. ISSN 1053-8119. doi: 10.1016/j.neuroimage.2019.05.060. URL https://www.sciencedirect.com/science/article/pii/S1053811919304525.

Steven J. Luck and Emily S. Kappenman. The Oxford Handbook of Event-Related Potential Components. Oxford University Press, December 2011. ISBN 9780199705870. Google-Books-ID: gItoAgAAQBAJ.

Shruti Naik, A. Banerjee, R. Bapi, G. Deco, and Dipanjan Roy. Metastability in senescence. 2017. doi: 10.1016/j.tics.2017.04.007. URL https://www.semanticscholar.org/paper/10672ce36487e27ccc53c548176411eddfa7ee86.

Amit Naskar, Anirudh Vattikonda, Gustavo Deco, Dipanjan Roy, and Arpan Banerjee. Multiscale dynamic mean field (mdmf) model relates resting-state brain dynamics with local cortical excitatory–inhibitory neurotransmitter homeostasis. Network Neuroscience, pages 1–26, 2021. ISSN 2472-1751. doi: 10.1162/netn_a_00197.

Anagh Pathak, Vivek Sharma, Dipanjan Roy, and Arpan Banerjee. Biophysical mechanism underlying compensatory preservation of neural synchrony over the adult lifespan. 5, June 2022. ISSN 2399-3642. doi: 10.1038/s42003-022-03489-4.

Julio Cesar Hernandez Pavon, Nils Schneider Garces, John Patrick Begnoche, Lee Miller, and Tommi Raij. TMS-EEG-MRI-fMRI-DWI data on paired associative stimulation and connectivity (Shirley Ryan AbilityLab, Chicago, IL), February 2022. URL https://openneuro.org/datasets/ds004024/versions/1.0.0.

Type: dataset.

Maria Concetta Pellicciari, Domenica Veniero, and Carlo Miniussi. Characterizing the cortical oscillatory response to tms pulse. Frontiers in Cellular Neuroscience, 11, 2017. ISSN 1662-5102. doi: 10.3389/fncel.2017.00038.

Ajay S. Pillai and Viktor Jirsa. Symmetry breaking in space-time hierarchies shapes brain dynamics and behavior. 2017. doi: 10.1016/j.neuron.2017.05.013. URL https://www.semanticscholar.org/paper/3a6809bc06f23d1312f5c03972ba261c42aa375f.

Malgorzata Plechawska-Wojcik, Monika Kaczorowska, and Dariusz Zapala. The artifact subspace reconstruction (ASR) for EEG signal correction. a comparative study. In Jerzy Świątek, Leszek Borzemski, and Zofia Wilimowska, editors, Information Systems Architecture and Technology: Proceedings of 39th International Conference on Information Systems Architecture and Technology – ISAT 2018, Advances in Intelligent Systems and Computing, pages 125–135, Cham, 2019. Springer International Publishing. ISBN 9783319999968. doi: 10.1007/978-3-319-99996-8_12.

Andreas Trier Poulsen, Andreas Pedroni, Nicolas Langer, and Lars Kai Hansen. Microstate EEGlab toolbox: An introductory guide, March 2018. Type: article.

Suman Saha, Priyanka Chakraborty, Amit Naskar, Dipanjan Roy, and Arpan Banerjee. Neuromolecular interactions guiding homeostatic mechanisms underlying healthy ageing: A view from computational microscope, 2023.

Ema Tanovic and Jutta Joormann. Anticipating the unknown: The stimulus-preceding negativity is enhanced by uncertain threat. International Journal of Psychophysiology, 139:68–73, May 2019. ISSN 0167-8760. doi: 10.1016/j.ijpsycho.2019.03.009. URL https://www.sciencedirect.com/science/article/pii/S0167876018311449.

Emmanuelle Tognoli and J. A. Scott Kelso. The metastable brain. Neuron, 81(1): 35–48, January 2014. ISSN 0896-6273. doi: 10.1016/j.neuron.2013.12.022. URL https://www.sciencedirect.com/science/article/pii/S0896627313011835.

N. Tzourio-Mazoyer, B. Landeau, D. Papathanassiou, F. Crivello, O. Etard, N. Delcroix, B. Mazoyer, and M. Joliot. Automated anatomical labeling of activations in spm using a macroscopic anatomical parcellation of the mni mri single-subject brain. 2002. doi: 10.1006/nimg.2001.0978. URL https://www.semanticscholar.org/paper/03d61a33796234b8bae5ac38de9b26c1c5ed9e2f.

Dimitri Van De Ville, Juliane Britz, and Christoph M. Michel. EEG microstate sequences in healthy humans at rest reveal scale-free dynamics. Proceedings of the National Academy of Sciences, 107(42):18179–18184, October 2010. doi: 10.1073/pnas.1007841107.

Irene Winkler, Stefan Haufe, and Michael Tangermann. Automatic classification of artifactual ICA-components for artifact removal in EEG signals. Behavioral and brain functions, 7:30, August 2011. ISSN 1744-9081. doi: 10.1186/1744-9081-7-30.

Wei Wu, Corey J. Keller, Nigel C. Rogasch, Parker Longwell, Emmanuel Shpigel, Camarin E. Rolle, and Amit Etkin. ARTIST: A fully automated artifact rejection algorithm for single-pulse TMS-EEG data. Human Brain Mapping, 39(4): 1607–1625, January 2018. ISSN 1065-9471. doi: 10.1002/hbm.23938. URL https://www.ncbi.nlm.nih.gov/pmc/articles/PMC6866546/.

