## Supplementary Results for "Metastability indexes global network effects post brain stimulation"

### Supplementary Material: Using metastability to identify the global changes in dynamic working point of the brain following brain stimulation

November 23, 2023

### 1 Supplementary Results

#### 1.1 Event Related Potential Analysis

Event Related Potentials (ERPs) are electrical changes that are time locked to sensory, motor or cognitive events (Picton et al., 2000). They are obtained through averaging activity around an event across a large number of trials. To understand preliminary results showing changes in metastability prior to the TMS pulse, an analysis of ERPs was also performed. Specifically, slow negative waves, which are documented signs of anticipation were tested for using established procedures (Luck and Kappenman, 2011). This analysis consisted of low pass filtering the data at 2 Hz, epoching around the pulse and then averaging across epochs and participants. 200 milliseconds preceding the TMS pulse were then visually inspected for a slow negative wave.

The ERP analysis revealed a Stimulus Preceding Negativity (SPN) with a central distribution prior to the TMS pulse in all 3 stimulation intensities (see Figure 1). The small magnitude of this effect could be explained by signal attenuation due to the aggressive preprocessing pipeline.

### 2 Supplementary Figures

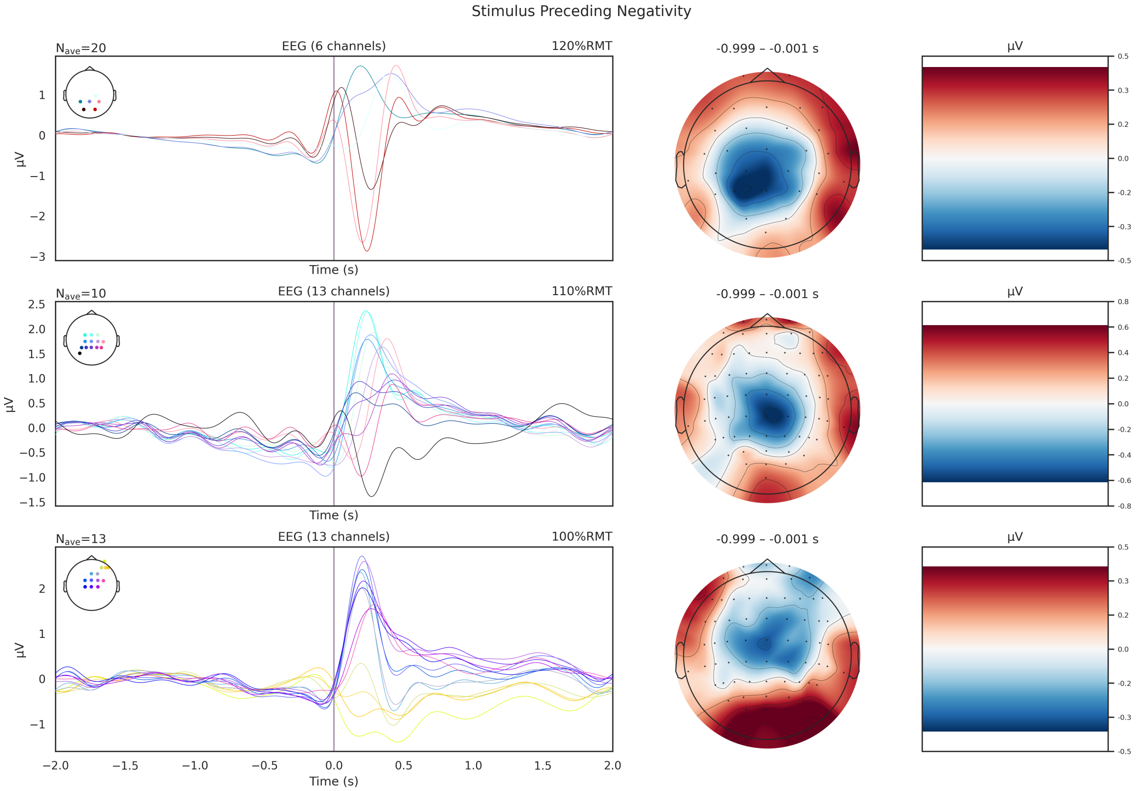


**Figure 1.** Stimulus Preceding Negativity (SPN) observed prior to the TMS pulse. Each row of plots pertains to a different stimulation intensity as indicated on the upper right of the ERP timeseries. The voltage distribution on the scalp is also shown indicating that the negativity is over the central electrodes.

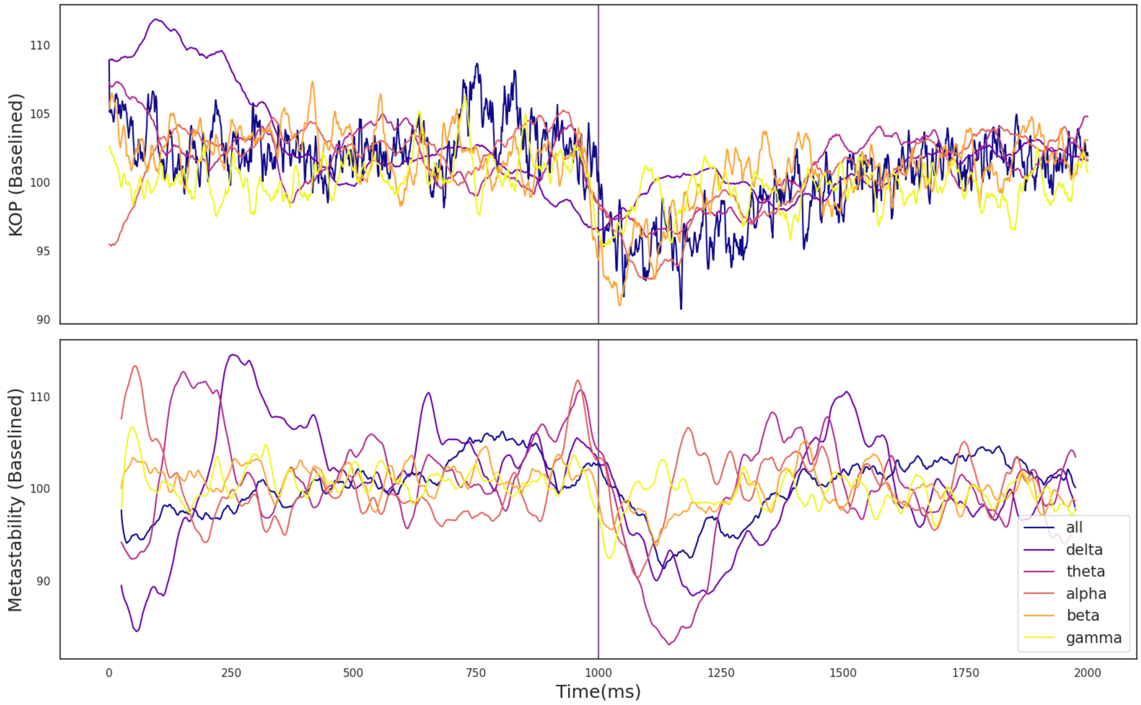


**Figure 2.** A) The effect of the TMS pulse on the Kuramoto Order Parameter. B) The effect of the TMS pulse on Metastability calculated in a sliding window. The TMS pulse is indicated by the vertical purple line. Both metastability and coherence are plotted as a percentage of a baseline value calculated as the mean between 525 and 1525 ms. Results are plotted in unique colors for each frequency band as per the legend. These results pertain to all electrodes and the 100% RMT stimulation condition.

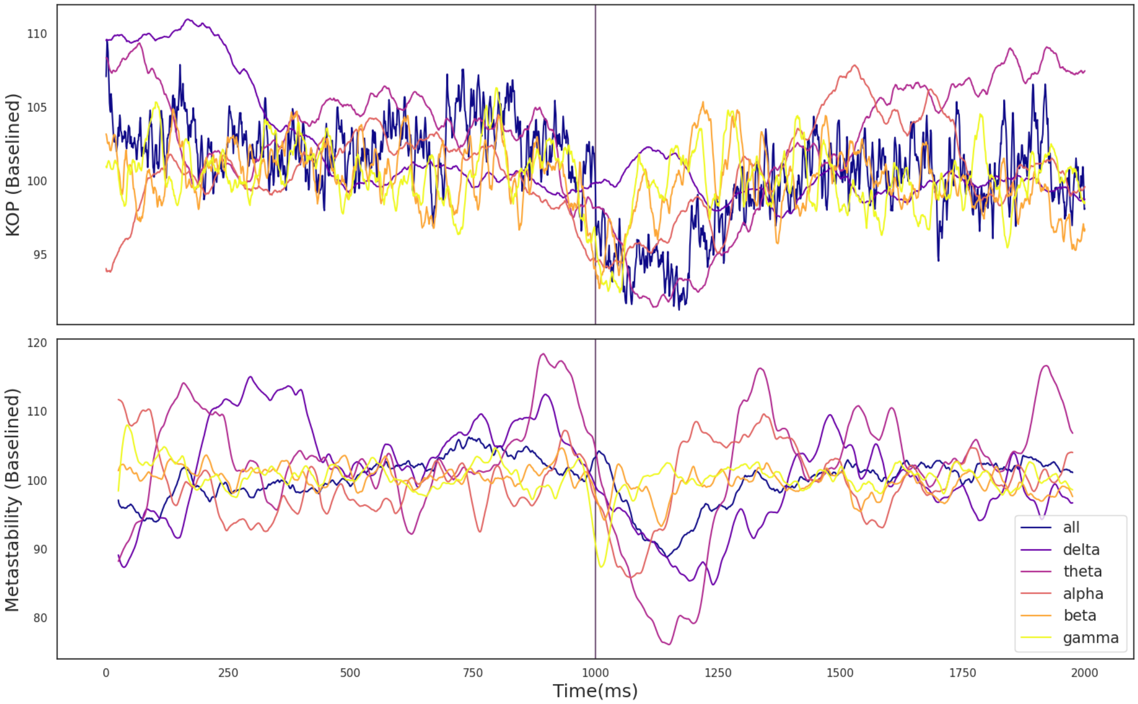


**Figure 3.** A) The effect of the TMS pulse on the Kuramoto Order Parameter. B) The effect of the TMS pulse on Metastability calculated in a sliding window. The TMS pulse is indicated by the vertical purple line. Both metastability and coherence are plotted as a percentage of a baseline value calculated as the mean between 525 and 1525 ms. Results are plotted in unique colors for each frequency band as per the legend. These results pertain to all electrodes and the 110% RMT stimulation condition.

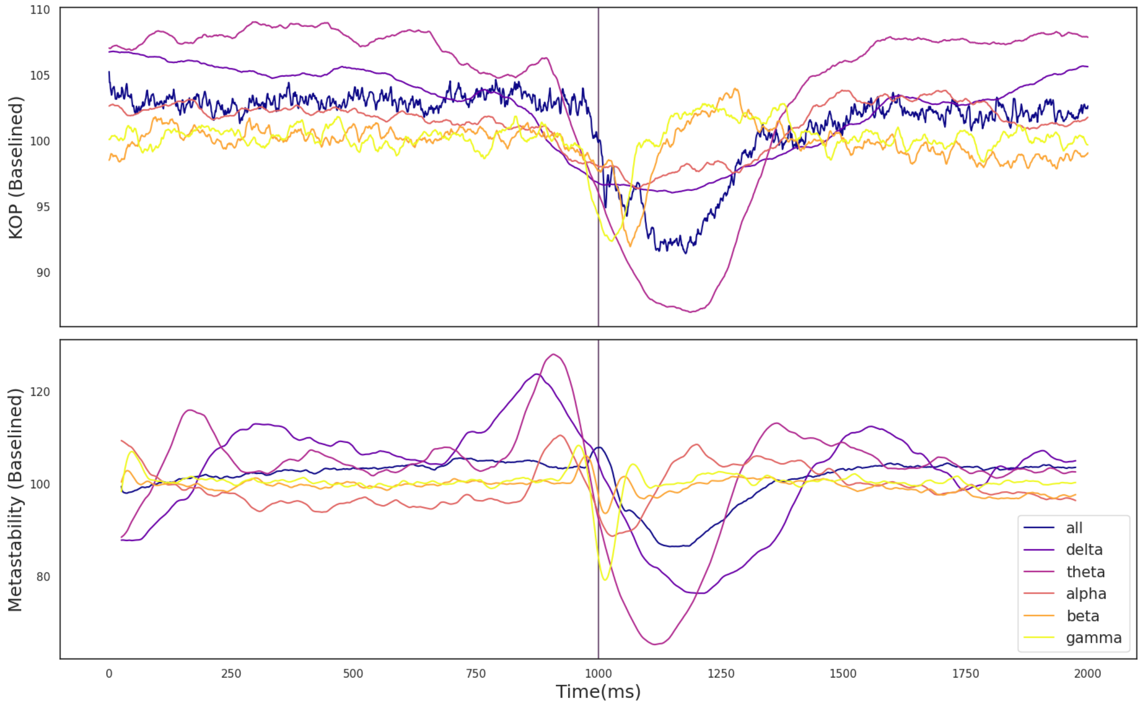


**Figure 4.** A) The effect of the TMS pulse on the Kuramoto Order Parameter. B) The effect of the TMS pulse on Metastability calculated in a sliding window. The TMS pulse is indicated by the vertical purple line. Both metastability and coherence are plotted as a percentage of a baseline value calculated as the mean between 525 and 1525 ms. Results are plotted in unique colors for each frequency band as per the legend. These results pertain to all electrodes and the 120% RMT stimulation condition.

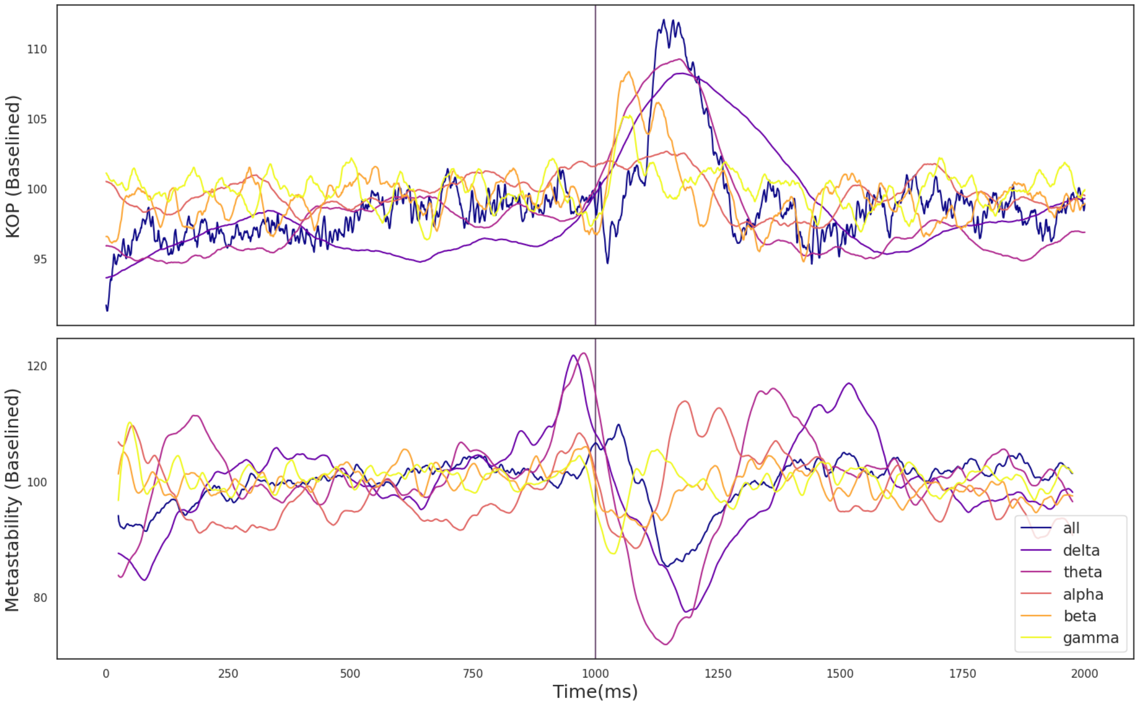


**Figure 5.** A) The effect of the TMS pulse on the Kuramoto Order Parameter. B) The effect of the TMS pulse on Metastability calculated in a sliding window. The TMS pulse is indicated by the vertical purple line. Both metastability and coherence are plotted as a percentage of a baseline value calculated as the mean between 525 and 1525 ms. Results are plotted in unique colors for each frequency band as per the legend. These results pertain to the fronto-central electrode group and the 100% RMT stimulation condition.

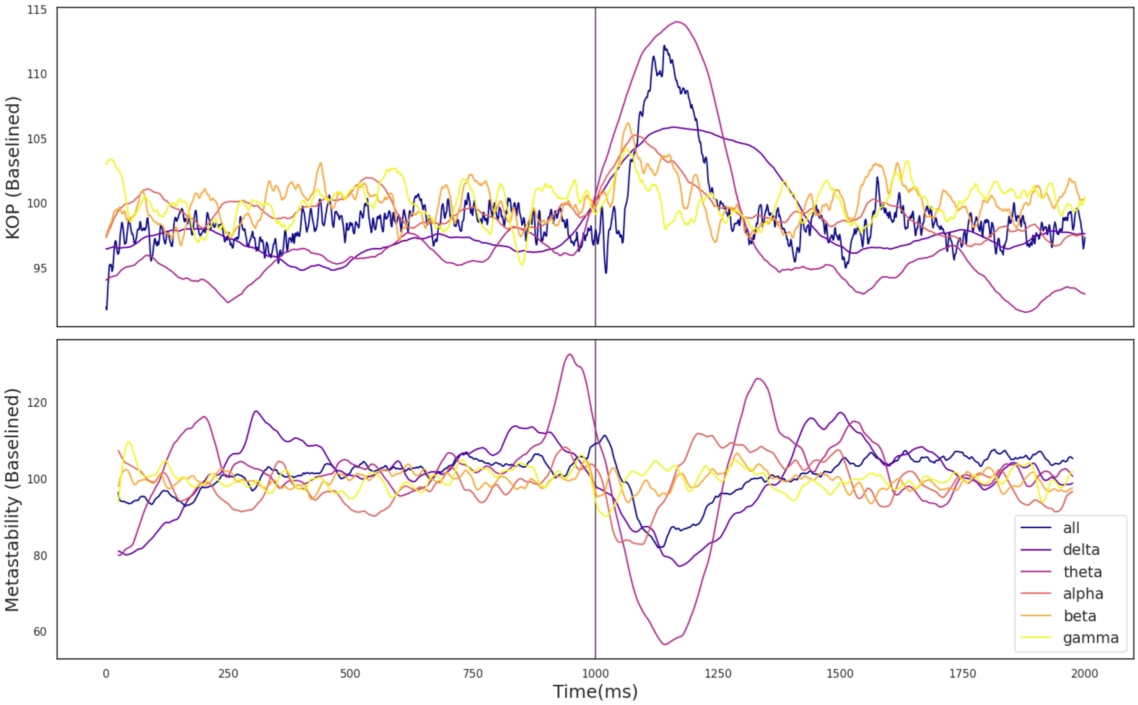


**Figure 6.** A) The effect of the TMS pulse on the Kuramoto Order Parameter. B) The effect of the TMS pulse on Metastability calculated in a sliding window. The TMS pulse is indicated by the vertical purple line. Both metastability and coherence are plotted as a percentage of a baseline value calculated as the mean between 525 and 1525 ms. Results are plotted in unique colors for each frequency band as per the legend. These results pertain to the fronto-central electrode group and the 110% RMT stimulation condition.

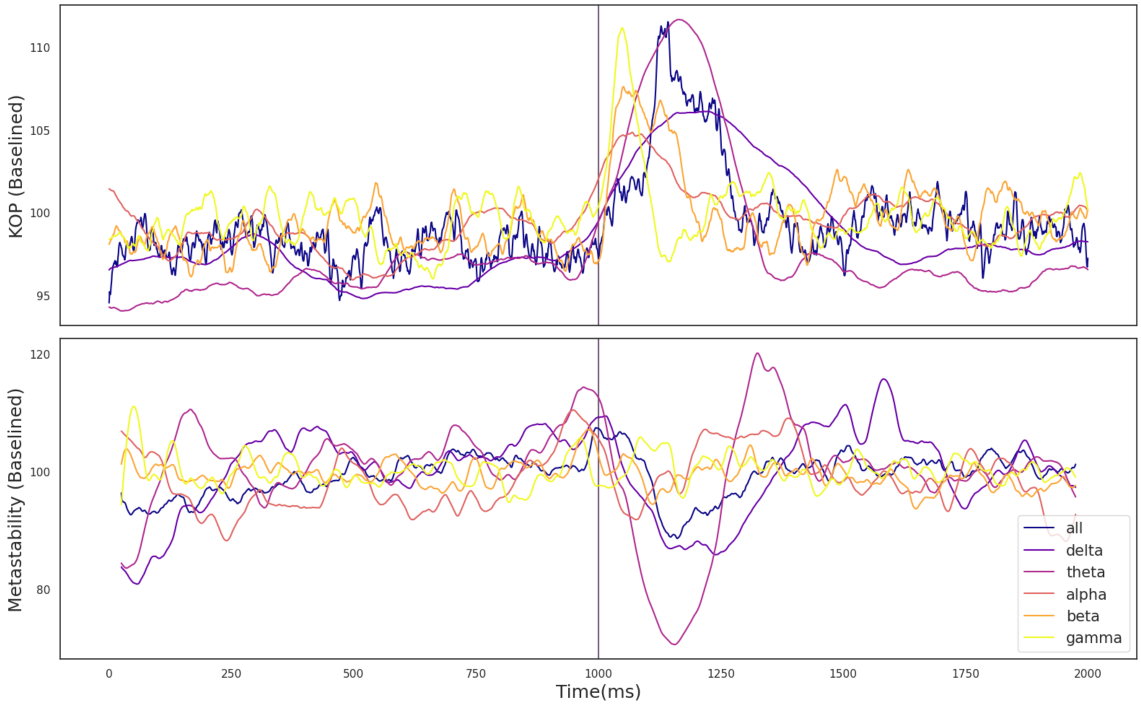


**Figure 7.** A) The effect of the TMS pulse on the Kuramoto Order Parameter. B) The effect of the TMS pulse on Metastability calculated in a sliding window. The TMS pulse is indicated by the vertical purple line. Both metastability and coherence are plotted as a percentage of a baseline value calculated as the mean between 525 and 1525 ms. Results are plotted in unique colors for each frequency band as per the legend. These results pertain to the temporo-occipital electrode group and the 100% RMT stimulation condition.

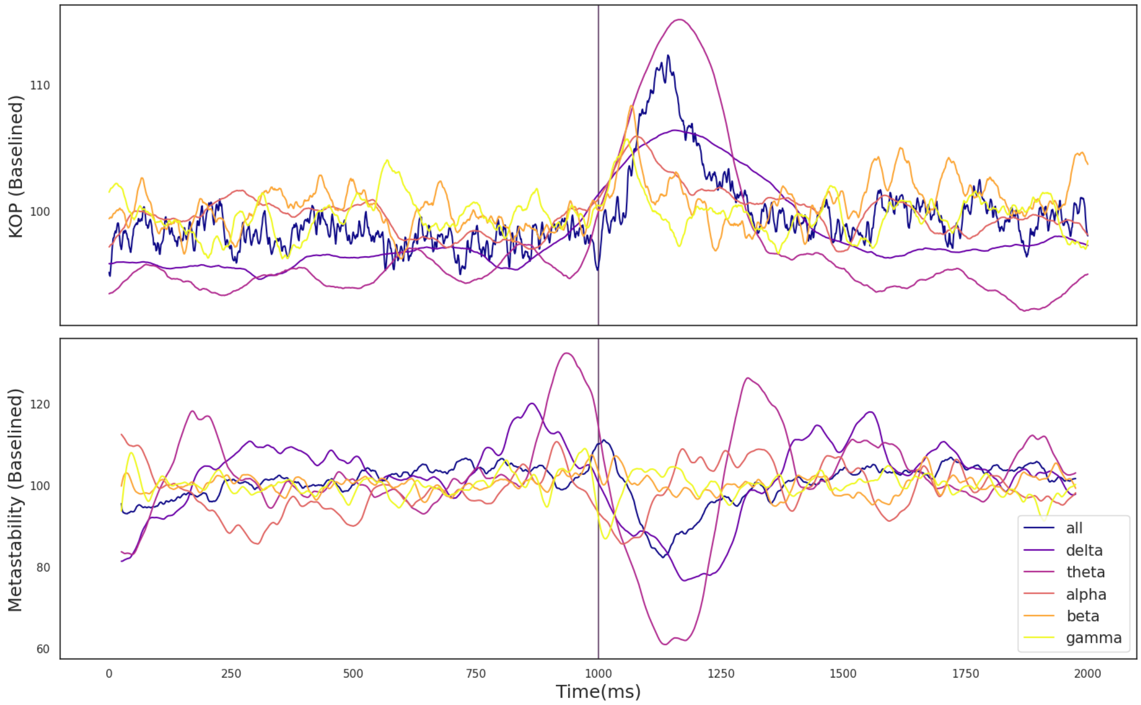


**Figure 8.** A) The effect of the TMS pulse on the Kuramoto Order Parameter. B) The effect of the TMS pulse on Metastability calculated in a sliding window. The TMS pulse is indicated by the vertical purple line. Both metastability and coherence are plotted as a percentage of a baseline value calculated as the mean between 525 and 1525 ms. Results are plotted in unique colors for each frequency band as per the legend. These results pertain to the temporo-occipital electrode group and the 110% RMT stimulation condition.

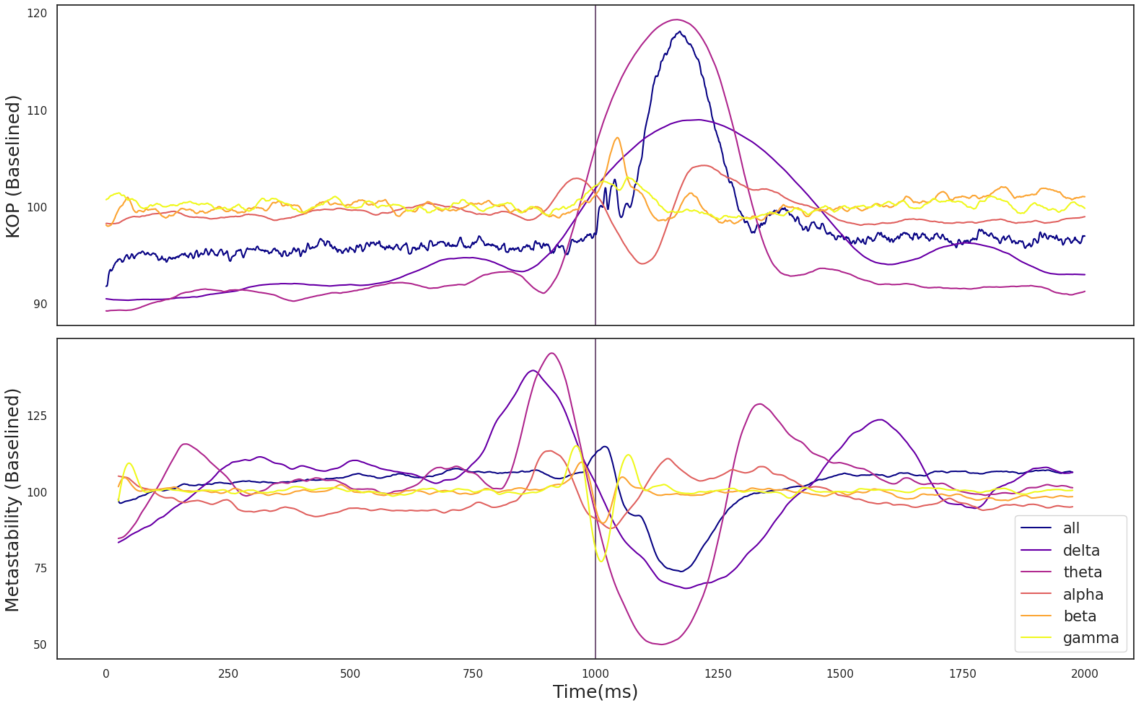


**Figure 9.** A) The effect of the TMS pulse on the Kuramoto Order Parameter. B) The effect of the TMS pulse on Metastability calculated in a sliding window. The TMS pulse is indicated by the vertical purple line. Both metastability and coherence are plotted as a percentage of a baseline value calculated as the mean between 525 and 1525 ms. Results are plotted in unique colors for each frequency band as per the legend. These results pertain to the temporo-occipital electrode group and the 120% RMT stimulation condition.

### References

Steven J. Luck and Emily S. Kappenman. *The Oxford Handbook of Event-Related Potential Components*. Oxford University Press, December 2011. ISBN 9780199705870. Google-Books-ID: gItoAgAAQBAJ.

T. Picton, S. Bentin, P. Berg, E. Donchin, S. Hillyard, Ray Johnson, Gregory A. Miller, W. Ritter, D. Ruchkin, M. Rugg, and Margot J. Taylor. Guidelines for using human event-related potentials to study cognition: recording standards and publication criteria. 2000. doi: 10.1111/1469-8986.3720127. URL https://www.semanticscholar.org/paper/4297e34b99c7a65917a139bec56e6325d6e3585d.
